# Local actuation of organoids by magnetic nanoparticles

**DOI:** 10.1101/2022.03.17.484696

**Authors:** Abdel Rahman Abdel Fattah, Niko Kolaitis, Katrien Van Daele, Andika Gregorius Rustandi, Adrian Ranga

## Abstract

Tissues take shape during development through a series of morphogenetic movements guided by local cell-scale forces. While current in vitro approaches subjecting tissues to homogenous stresses, it is currently no possible to recapitulate highly local spatially varying forces. Here we develop a method for local actuation of organoids using embedded magnetic nanoparticles. Sequential aggregation of magnetically labelled human pluripotent stem cells followed by actuation by a magnetic field produces localized magnetic clusters within the organoid. These clusters impose local mechanical forces on the surrounding tissue in response to applied global magnetic fields. We show that precise, spatially defined actuation provides short-term mechanical tissue perturbations as well as long-term cytoskeleton remodeling. We demonstrate that local magnetically-driven actuation guides asymmetric growth and proliferation, leading to enhanced patterning in human neural organoids. We show that this approach is applicable to other model systems by observing polarized patterning in paraxial mesoderm organoids upon local magnetic actuation. This versatile approach allows for local, controllable mechanical actuation in multicellular constructs, and is widely applicable to interrogate the role of local mechanotransduction in developmental and disease model systems.

## Introduction

During embryogenesis, tissues are generated by morphogenic events which are tightly controlled by temporally defined mechanical forces. Forces can be generated by intrinsic cellular contractility or as a consequence of adjacent tissues interacting with each other^1,2^. Cells respond to such forces through the activation of mechanosensitive pathways^3,4^, thereby generating mechano-responsive feedback which serve to change their shape, proliferation and fate specification^5^. In particular, intrinsic forces generated by the tissue undergoing morphogenesis play an important tissue-shaping role. For example during the developing early central nervous system, the formation of the neural tube has been attributed to neural plate folding without assistance by surrounding tissues^6,7^. Indeed, the initiation of neural plate folding has been attributed to localized apical constriction at the median hinge point^8^ that later controls floor plate cell shape and cell cycle^9,10^. While it is well established that the response of cellular subpopulations within a tissue to local forces is necessary for morphogenic programs to engage in the development of complex tissue architecture, how such forces generate local sub-tissue level responses and how such responses coordinate tissue-wide growth and patterning effects remains unclear.

Organoids are ideal model systems to address such questions as they reflect key elements of the cellular diversity and architectural complexity seen in-vivo while allowing control over the mechanical and biochemical microenvironments which help direct differentiation and patterning. Synthetic extracellular matrices such as polyethylene glycol (PEG) hydrogels have shown how the matrix stiffness modulation could not only sustain intestinal stem cell growth and expansion^11^, but could also guide organoid differentiation^12^. The ability to tailor the mechanics such microenvironments has been used to define a matrix stiffness range which could specify cell patterning in mouse neural tube organoids^13^. Such studies rely primarily on predefined gel stiffnesses or matrix degradation, where microenvironment mechanical characteristics cannot be actively modified, thereby missing the recapitulation of dynamic mechanical events seen in-vivo. To impose active mechanical perturbations to organoid cultures, we recently developed a mechanical device which could control force modulation on organoids, and demonstrated an enhancement of floor plate patterning in organoids upon actuation^14^. This approach can only impose external and largely homogeneous forces on tissues, allowing an understanding of mechanoregulatory effects of global forces but not local ones.

Various approaches have been implemented to mechanically stimulate tissues locally. Optical tweezing can deliver subcellular point forces but is limited to piconewton ranges and thus to applications required only weak forces, such as the mechanical actuation of ion channels in neurons^15^. Higher forces can be delivered using techniques such as microindentation^16^ and micro-aspiration, which has been used in a variety of applications such as in elucidating electrical-mechanical coupling in retinal cells^17^ or in uncovering a biophysical mechanism in the regeneration of mucociliated epithelium^18^. While able to deliver local forces, such techniques require tissue-surface contact and are thus limited to stimulate local but external forces. In 3-dimentional developing tissues, force generation is not limited to the surface and therefore the ability to provide internal mechanical stimulation is needed.

To mechanically stimulate tissues internally, magnetic nanoparticles (MNPs) have been used, whereby, once embedded in a tissue, they can respond to a magnetic field and cause localized mechanical stresses. For example ferrofluids embedded in zebrafish embryos were used to demonstrate a posterior decrease in tissue stiffness^19^. This technique is suited for in-vivo investigations, however due to the size of the magnetic actuators (∼100 µm), may be less adequate for smaller in-vitro model systems. MNP internalization offers an alternative that allows for cell-level in-vitro actuation. Internalized MNPs have been used to demonstrate polarized increases in cytoskeleton rearrangement^20^ in response to mechanical stresses from an external magnetic field. This type of remote actuation is scalable to multicellular constructs, for example when applied to mouse embryonic stem cell aggregates to facilitate cardiac differentiation though cyclic mechanical forces produced by a static magnet^21^.

While demonstrating the power of magnetic, remote, and local actuation in-vitro, these studies have been limited to inducing local forces in 2D or imposing global forces in 3D. There has thus far not been any way to impose more in-vivo relevant local forces within organoids, and by extension the corresponding mechanoregulatory effects remain largely underexplored. Here, we develop an approach to actuate local regions within an organoid using embedded magnetic nanoparticles, which we term magnetoids and show that global magnetic fields can be used to generate internal and local forces, modulating growth, morphogenesis and patterning within the organoid body.

### Magnetoid generation

Our approach aimed at providing localized forces by concentrating magnetic nanoparticle into clusters (MagCs) within a region of an organoid. To create a magnetoid, we developed a simple protocol which ensured that under a magnetic field, only the tissue in close proximity to the MagC would be mechanically actuated in response to the magnetic field, thus providing local mechanical forces.

In order to ensure that the location of the MagC could be visualized by fluorescence microscopy, we first labelled MagCs with carboxylated fluorescent particles (FPs) (**Fig.1a**). The fluorescent MagCs were then incubated with single hPSCs to magnetize the cells. For maximal cell magnetization, a high concentration of NPs is desirable, however this may be deleterious to cell viability^22–26^. To optimize the concentration of MNPs, we incubated hPSCs with a range of MNP concentrations, and determined that 1,000 μg/ml was the highest MNP concentration which would maintain a high cell viability (**Fig.1b**). In contrast with most previous uses of MNPs, here our MNP clusters do not undergo uptake through endocytosis due to their size (∼2 μm) (**supplementary Fig.1a**), and are instead adsorbed on the membrane exterior and found primarily on top (apical side) of the hPSC colonies, as demonstrated by colocalization with the expression of ZO1 (**supplementary Fig.1b**).

**Figure 1:**
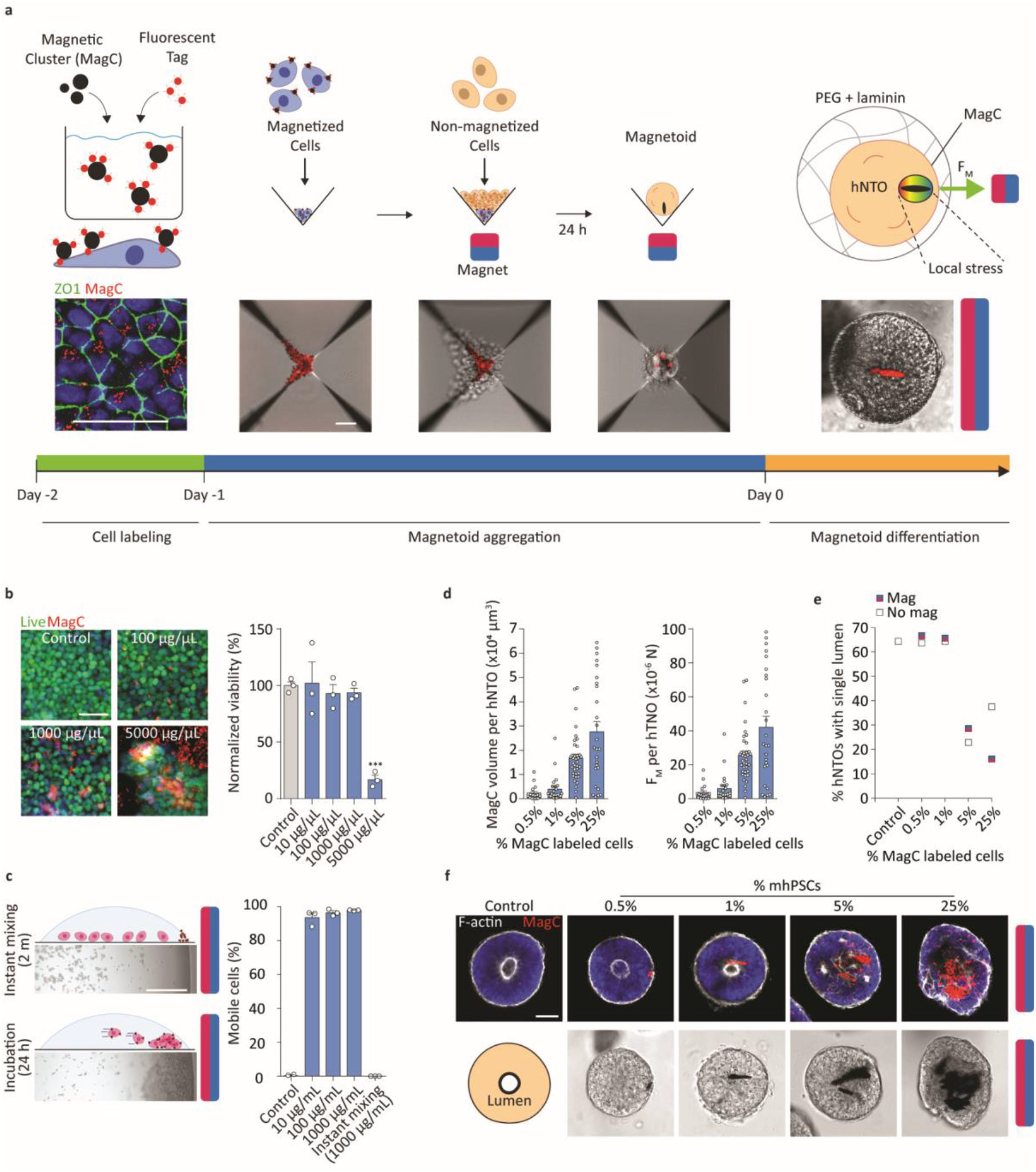
Magnetoid generation protocol. **a** Magnetoid generation and hNTmO differentiation protocol. Scalebars 50 μm. **b** Toxicity test with normalized hPSC count after 3-day exposure to various concentrations of MagCs (n = 3). Representative images showing live hPSCs in control and MagC conditions (100, 1,000, and 5,000 μg/mL) (n = 3). Scalebar 50 μm. **c** Representative images showing mhPSC magnetic attraction (n = 2). Percentage of mobile cells at various MagC concentrations (n = 3). Scalebar 500 μm. **d** Estimation of MagC volumes in day 11 hNTmOs at various mhPSC proportions (n = 3 for >20 magnetoid per condition). Estimation of force generation in day 11 hNTmOs at various mhPSC proportions (n = 3 for >20 magnetoid per condition). **e** Average lumen formation in day 11 hNTmOs at various conditions and with various mhPSC proportions (n = 2 for >30 magnetoid per condition).**f** Representative images showing F-actin expressions in day 11 hNTmOs at various mhPSC proportions (n = 3). Scalebar 50 μm. Error bars SEM.

We next verified that the MagCs adhered to the cells to produce magnetized hPSC (mhPSCs), and found that the adhesion was strong enough to guide cells by magnetophoresis in a liquid medium where nearly 100% of cells display mobility, i.e. cells were successfully magnetized (**Fig.1c**). Of note, the incubation of MagCs with cells was necessary since instant mixing of MagCs with single hPSCs did not provide adequate membrane adsorption of the particles (**Fig.1c**). To estimate the volume of adsorbed MagCs per cell, a small static magnet was used to move single mhPSCs in suspension in a growth medium. MagC volume was found to be ∼7.3 ×10^3^ μm^3^ per cell by equating Stoke’s drag force to the magnetic force for cells undergoing magnetophoresis (**see methods, supplementary Fig.2**).

Magnetized pluripotent aggregates with localized MagCs (**Fig.1a**) were then produced by mixing magnetized and non-magnetized hPSCs through consecutive centrifugation steps, followed by a 1 day incubation under a magnetic field. These magnetized pluripotent aggregates were then embedded in PEG hydrogels with a stiffness of 2 kPa as previously described^14^, and could then be differentiated under constant or periodic magnetic actuation to impose internal local forces.

To investigate the forces generated by the MagCs, we generated human neural tube magnetoids (hNTmOs) following a previously described human neural tube differentiation protocol^14^. We first evaluated the magnetic field produced by a static magnet placed adjacent to a standard well-plate. We found that magnetoids situated furthest from the magnet surface were subjected to the lowest forces (**See methods, supplementary Fig.3a**). While these low forces led to MagC fragmentation, higher forces closer to the magnet led to high MagC aspect ratios and maintained cluster localization (**supplementary Fig.3b-c**). Thus to maintain consistent force localization and highest force values in our investigation, magnetoids were placed in wells closest to the magnet surface.

We next investigated the effect of mhPSC proportion on force generation. We found that increasing mhPSC proportion increased MagC volume and by extension the resultant force (**Fig.1d**). To ensure the embedded MagCs do not disrupt the magnetoid cytoarchitecture, an important consideration for correct differentiation and growth^13,14^, we investigated the cytoskeleton arrangement in day 11 hNTmOs with varying mhPSC proportions (**Fig.1e-f**). On average, low MagC volumes of ∼5.7×10^3^ μm^3^ correlated with single lumen magnetoids indicating correct apicobasal polarization (**supplementary Fig.4**). Indeed, only the low mhPSC proportions (≤1%) that produce small MagC volumes resulted in single lumen magnetoids with the same frequency as the MagC-free control organoids. In contrast, mhPSC proportions >1% reduced single lumen formation frequency regardless of magnetic field application (**Fig.1e-f**). Thus while 5 and 25% mhPSCs-magnetoids generated more force, 1% mhPSCs was chosen in our study as it provided the maximum force (∼6.0×10^−6^ N) with minimal impact on organoid cytoskeletal organization. This force was also found to be in the range of those previously reported to stall neural tube folding in amphibia^27^, and suggest that the magnetoid model system is able to attain relevant in-vivo forces.

Overall, magnetoids can be created with localized MagCs that are responsive to an external magnetic field while maintaining organoid cytoarchitecture. While forces can be simply generated by a static magnet, their strength and mode will likely affect the MagC-magnetoid interactions. We therefore thought to explore various force profiles to evaluate such interactions.

### Short and long term magnetic actuation in magnetoids

The temporal dynamics of force applications are important in studying the mechanobiology of cellular processes, which range from seconds and minutes, such as in heart contractions, or days for developmental events such as neural tube morphogenesis. We therefore thought to investigate the applicability of the magnetoid model system in emulating various force dynamics regimes. We evaluated the magnetoids response to magnetic actuation by observing tissue displacement and cytoskeleton rearrangement.

To evaluate the short-term response of organoids to fast-changing forces, an actuation burst was provided by introducing a magnetic field to a magnetoid, which was observed over 30 seconds, followed by magnet removal. This short perturbation event is far below reported force application durations (∼1 h) needed for cytoskeletal reorganization^28^. Upon magnetic field exposure, the embedded MagCs attempt to align their long axis with the magnetic field lines, pushing against the surrounding tissue and providing actuation (**Fig.2a-b**). Since small variations occur in the displacement of different MagCs when subjected to a magnetic field, for comparison purposes we normalized the tissue displacement map to the distance travelled by the corresponding MagC, d_M_, upon actuation. We found that tissues positioned >25 d_M_ from the MagC undergo minimal displacement (**Fig.2c**). Since the size of magnetoids are on the order of ∼100 d_M_ these results suggest that magnetoids can be used to deliver fast-changing localized forces actuating only the tissue in close proximity, leaving the remainder of the magnetoid unaffected.

**Figure 2:**
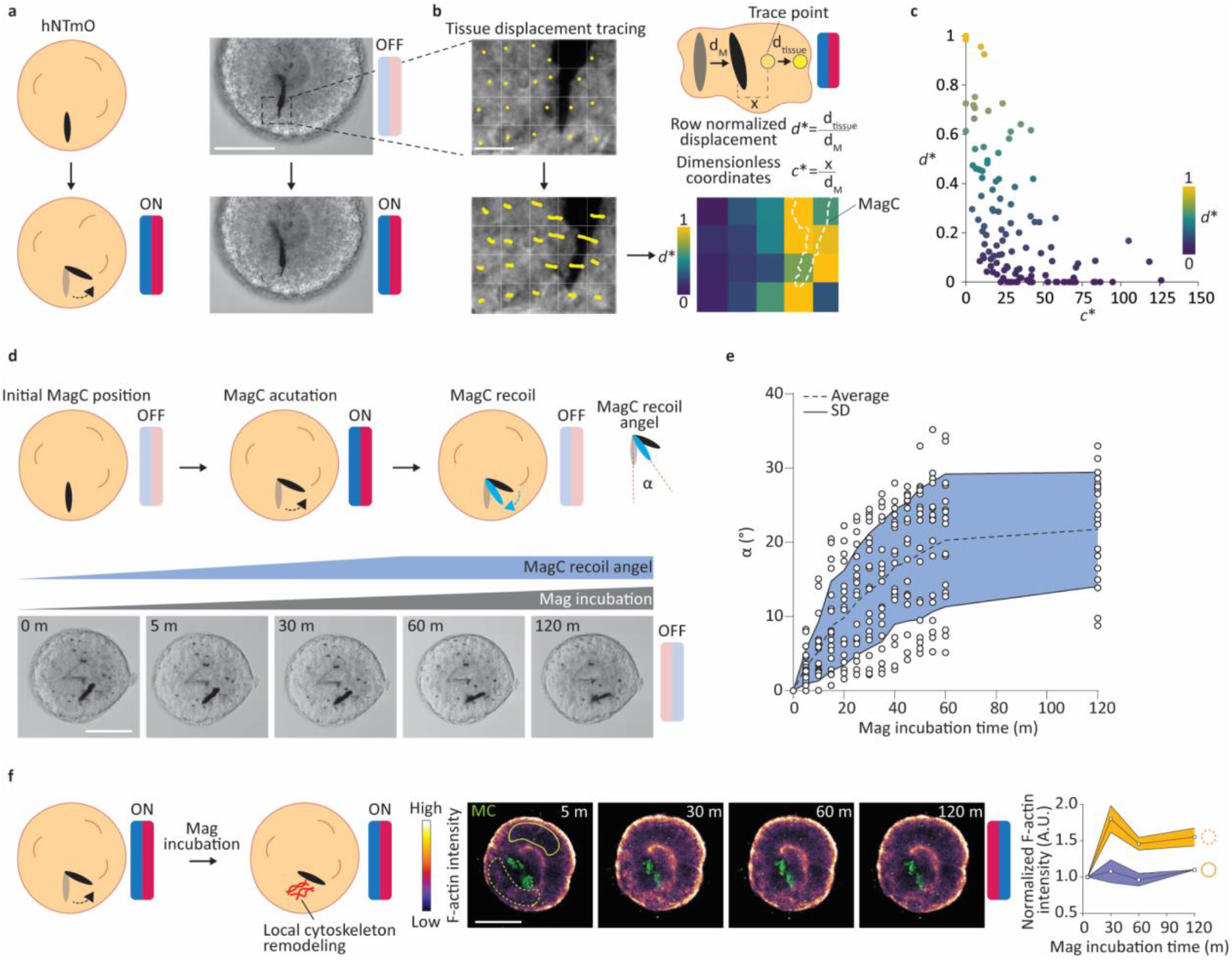
Short- and long-term local magnetic actuation. **a** Schematic representation of local short-term actuation of MagCs. Representative images of day 11-hNTmOs at with and without the influence of magnetic field (n = 2 for a total of 15 hNTmOs). Scalebar 50 μm. **b** Schematic representation of tissue displacement. Representative images of tissue tracing measurements and accompanied heatmap (n = 2 for a total of 15 hNTmOs) Scalebar 10 μm. **c** Correlation of dimensionless tissue and MagC displacements (n = 2 for a total of 15 hNTmOs). **d** Schematic representation of MagC actuation and recoil. Representative images of MagC actuation and recoil during long-term magnetic incubation (n = 3 for >15 hNTmOs). Scalebar 50 μm. **e** Recoil angle α as a function of magnetic incubation time (n = 3 for >15 hNTmOs). Dashed line represents the average α values and solid lines represent SD. **f** Schematic representation for long-term magnetic incubation and local cytoskeleton remodeling. Representative images showing F-actin expression of the same hNTmO at various magnetic incubation times and accompanied F-actin intensity close to and far away from the MagCs (n = 2 for a total of 5 hNTmOs). Solid line represents average and colored boundaries represent SD. Scalebar 50 μm.

Slow-changing forces can also be generated by subjecting magnetoids to constant magnetic fields for longer time periods. In order to investigate the transition from fast (bursts) to slow (constant) forces, we evaluated the recoil angle, α, of MagCs after removal of the magnetic field (**Fig.2d**). Short actuation times led MagCs to recoil to their original orientation once the magnetic field was lifted (**Fig.2e**). In contrast, longer actuation under a magnetic field led to higher values of α that stabilized after 1 h. These results suggest that constant forces applied over longer durations likely trigger cytoskeletal rearrangements that reinforce new MagC positions while preventing their return to their original position upon magnet removal (**Fig.2e**). Indeed, using an F-actin reporter hPSC line, we show that constant actuation is accompanied by cytoskeletal rearrangements and only in close proximity to the MagCs (**Fig.2f**).

These results suggest that mechanical memory of the initial MagC position is retained through the cytoskeletal architecture. While a short burst of magnetic actuation allows for the application of fast-changing forces while retaining mechanical memory, longer, constant forces appear to update this mechanical information by remodeling the local cytoskeletal architecture. Such cytoskeletal modulations play a particularly important role in neural development^8,13,14,29^. Therefore, we used the neural differentiation of magnetoids under constant magnetic actuation as a model system to investigate the effects of local actuation on neural tube morphogenesis and patterning.

### Local magnetic actuation biases growth and proliferation in magnetoids

Neural tube development relies on differential cell cycle regulation and growth^30^ where floor plate cells^9,10^ proliferate less than intermediate domains^31^. To investigate whether differential forces can guide these processes, we first investigated morphological changes in hNTmOs. Organoids without applied magnetic field, with or without MagCs, were seen to grow symmetrically with an average P_m_/P_a_ ∼1.05 and ∼1.01 respectively (**Fig.3a**), where P_m_ and P_a_ are the measured linear growth toward and away from the magnet respectively. In contrast, constant magnetic actuation over 11 days biased growth away from the magnet (P_m_/P_a_ ∼0.61). Despite this growth bias, we observed no significant differences in hNTmO sizes between all three conditions (**Supplementary Fig.5**), suggesting that magnetic forces do not change the overall proliferation capacity of organoids but only bias them directionally. To determine whether the observed growth bias was due to differential proliferation or internal deformation of the hNTmOs, we performed EdU staining. Proliferation appeared largely similar in the absence of a magnetic field, where the ratio of EdU intensity between the regions facing the magnet (EdU_m_), and those facing away from the magnet (EdU_a_), resulted in a EdU_m_/EdU_a_ ∼1.28 and ∼1.55 in magnetoids with and without MagCs respectively (**Fig.3b**). However, application of the magnetic field produced a significant proliferation bias away from the magnet with EdU_m_/EdU_a_ ∼0.80. These results suggest that after the initial realignment of the MagCs under magnetic incubation conditions, causing reorganization of the actin cytoskeleton (**Fig.2b** and **c**), the subsequent effect of the sustained magnetic force is to modulate growth and proliferation. Indeed, MagCs likely compress the tissue facing the magnet thereby counteracting growth in this region, while further away from the MagC position, unrestricted tissue can freely proliferate. Local actuation can therefore modulate local cell cycle to bias global growth direction, suggesting that bending of the neural plate in-vivo at the median hinge point can mechanically restrict growth and provide the necessary bias for intermediate domain proliferation.

**Figure 3:**
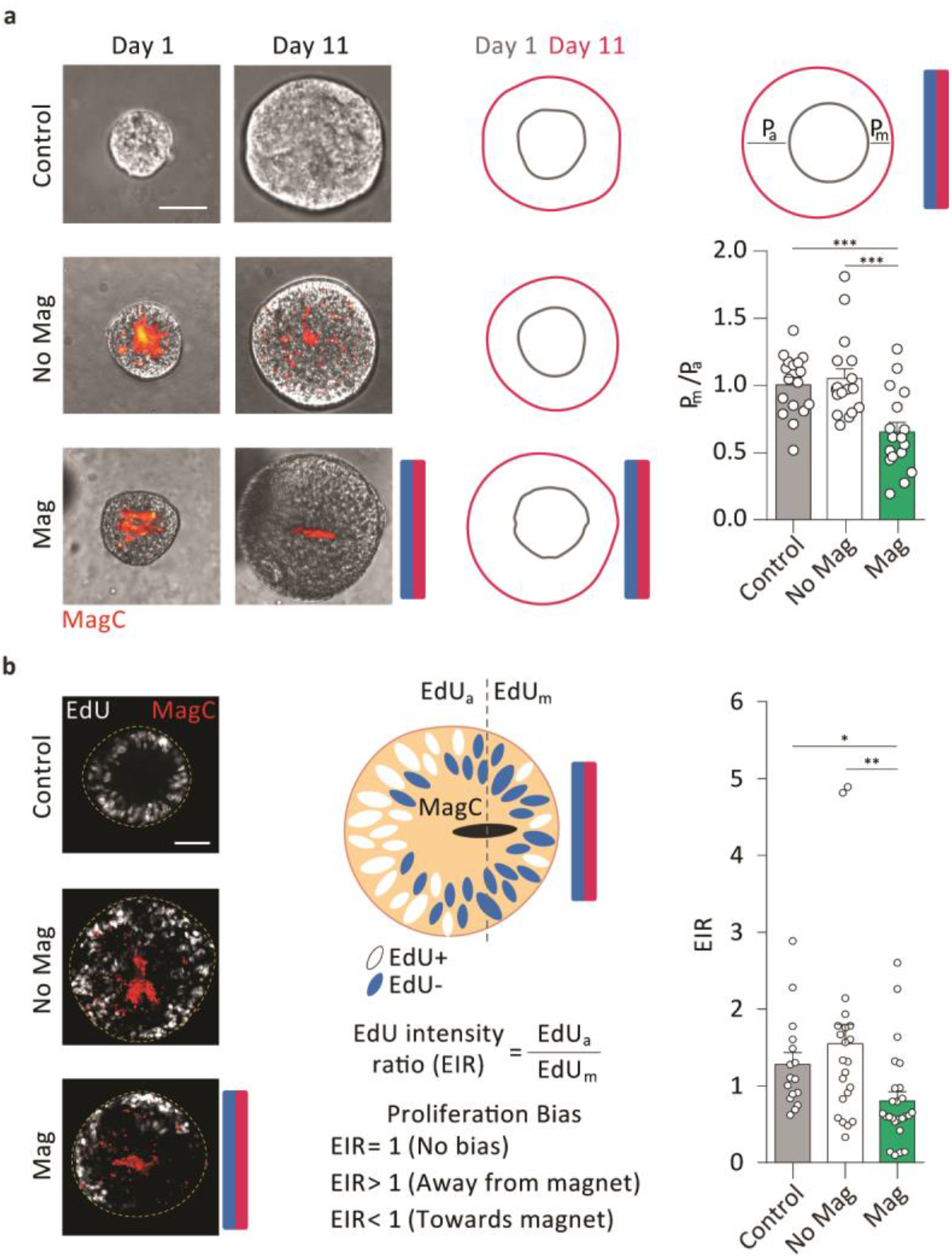
Local magnetic actuation biases growth and proliferation. **a** Representative images of day 1 and day 11 hNTmOs under various actuation conditions (n = 3). Schematic representation of day 1 and day 11 hNTmO contours. Growth ratio of hNTmOs for various actuation conditions (n = 3 for >15 hNTmOs pre condition). Scalebar 50 μm. **b** Representative images for EdU staining of day 11 hNTmOs under various actuation conditions (n = 2). Schematic representation of biased proliferation in hNTmO under the influence of magnetic field. Proliferation ratio of day 11 hNTmOs for various actuation conditions (n = 2 for >15 hNTmOs per condition). Scalebar 50 μm. Error bars SEM.

### Local magnetic actuation guides tissue-wide patterning in magnetoids

Mechanical stresses have been previously shown to guide cell fate specification and patterning in mouse^13,32^ and human^14^ NTOs. We therefore thought to investigate domain specification in our hNTmOs to evaluate mechanically-driven bias in patterning. We focused our analysis on two differentially regulated and mutually exclusive regions in the neural tube, the floor plate and intermediate domains. Floor plate induction, as marked by the transcription factor FOXA2, remained unchanged for hNTmOs with or without magnetic actuation when compared to the control (**Fig.4a**). Upon magnetic actuation, floor plate patterning frequency increased to ∼60% of FOXA2+ hNTmOs compared to ∼30% in control and hNTmOs not subjected to a magnetic field. Similarly, while intermediate fate induction, marked by PAX6, remained consistent across control organoids and hNTmOs regardless of actuation, patterning of the intermediate domain increased to ∼80% upon magnetic actuation (**Fig.4b**). We next thought to investigate whether cell fate was biased away or towards the magnet by evaluating the average FOXA2 or PAX6 fluorescence intensities away and towards the magnet as in the EdU case (**See methods, Fig.4c**). FOXA2 cells were found to pattern towards the magnet with a FOXA2_m_/FOXA2_a_ ∼1.30 in actuated magnetoids compared to a value of ∼1.03 and ∼1.02 for control organoids and unactuated magnetoids respectively (**Fig.4c**). In contrast, PAX6 cells were present in higher proportion away from the magnet in magnetoids with a PAX6_m_/PAX6_a_ ∼0.86 compared to a value of ∼1.01 and ∼1.00 for control organoids and unactuated magnetoids respectively (**Fig.4c**). Linking these results to observations of proliferation suggests that floor plate cells pattern in regions with growth restriction and lower proliferation while those of the intermediate fates are preferentially found in regions with higher proliferation. These observations are in line with in-vivo studies^31^ showing that the median hinge point comprising of floor plate cells is a region of low cell proliferation, while the intermediate domain between dorsal and ventral regions proliferates rapidly during the course of neural tube development, suggesting that cell proliferation may be due to less mechanical restriction.

**Figure 4:**
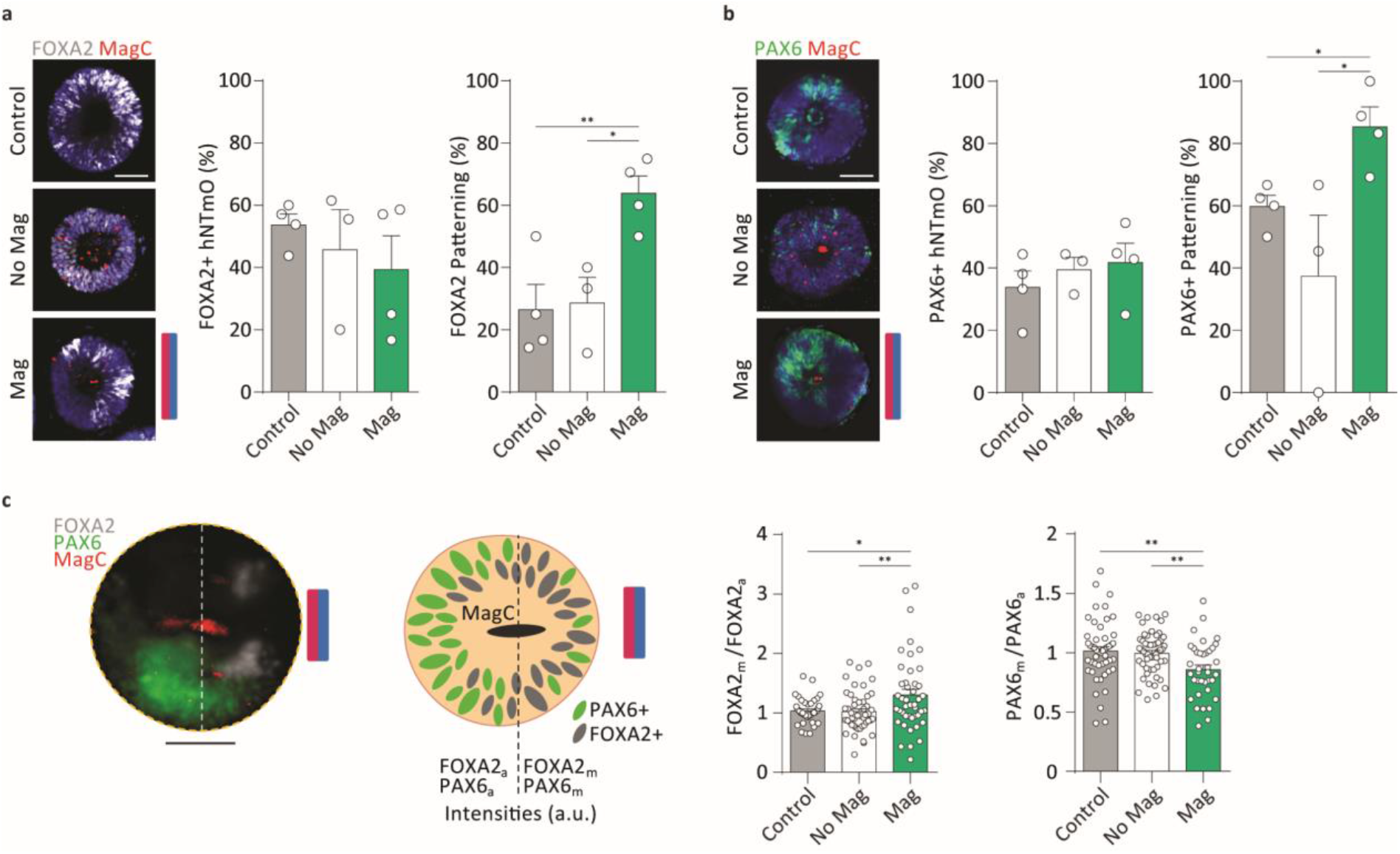
Local magnetic actuation biases patterning in hNTmOs. **a** Representative images showing FOXA2 expression in day 11 hNTmOs under various actuation conditions. Quantification of FOXA2 induction and patterning in hNTmOs under various actuation condition (n = 4 except for the No Mag condition n = 3 for a total of >60 hNTmOs for all data points per condition). Scalebar 50 μm. **b** Quantification of PAX6 induction and patterning in hNTmOs under various actuation condition (n = 4 except for the No Mag condition n = 3 for a total of >60 hNTmOs for all data points per condition). Scalebar 50 μm. **c** Representative image of day 11 hNTmO under magnetic actuation showing biased expression of FOXA2 and PAX6 (n = 4). Schematic representation of biased FOXA2 and PAX6 expressions under magnetic actuation condition. Quantification of FOXA2 intensity ration for day 11 hNTmO under various actuation conditions (n = 4 for a total of >40 hNTmO per condition). Scalebar 50 μm. Error bars SEM.

Interestingly, as the neural tube increases in size, cell patterning is maintained through antiparallel gradients operating across the growing tissue with cells interpreting ranges of biochemical cues at various distances^33^. With regards to these observations, we demonstrated a mechanical analogue where the distance between FOXA2+ cells and the MagC correlated with different ranges of magnetic forces, and where larger forces allowed for more distant expressions of FOXA2 from the MagCs (**Supplementary Fig.6**). A similar correlation was observed with PAX6+ cells, in regions furthest from the magnet (**Supplementary Fig.6**). The results suggest that the position of cells in relation to the source of the mechanical force is involved in specify its fate, and may serve as a mechanism for pattern regulation mechanisms across growing tissues.

To demonstrate the wider applicability of this local magnetic actuation system to other in-vitro model systems, we applied our approach to organoids differentiated towards paraxial mesoderm identity. Over the 8 days of differentiation, a gradual increase in PAX3 expression was observed in human paraxial mesoderm magnetoids (**Supplementary Fig. 7a**), corresponding to the differentiation trajectory of paraxial mesoderm cells in-vivo^34^. These changes in cell identity were coupled to lower proliferation^34^. Since actuated hNTmOs exhibited proliferation bias, we thought to investigate PAX3 expression in paraxial mesoderm magnetoids. As for FOXA2 fate in hNTmOs, PAX3 expression in actuated paraxial mesoderm magnetoids was biased towards the magnet surface (**Supplementary Fig. 7b**), suggesting that mechanical restriction caused by the magnetic force guided PAX3 polarization towards regions of lower proliferation, reflecting in-vivo observations.

Altogether, our results demonstrate that internal local tissue actuation can guide tissue-wide organoid morphology, growth, and patterning. These local mechanical forces are achieved using a simple protocol to embed local magnetic tissue actuators in organoids. We show that this approach can generate both fast or slow-changing forces, and that rapid short-term actuation deforms local tissue, while longer-term actuation triggers cytoskeleton rearrangement processes and tissue-scale differential growth and cytoarchitecture remodeling. We also demonstrate local patterning enhancement in neural tube and paraxial mesoderm organoids, suggesting a broader role of actuation in enhancing patterning. While no preferential direction was previously observed in neural tissue under exogenous actuation^14,35^, our results demonstrates that local actuation guides directional patterning bias. Moreover, while we demonstrate the use of static magnets for actuation, it is conceivable to stimulate the magnetoids through an large number of magnetic configurations stemming from static or electro-magnets, providing design freedom for the user to fit the needs of the specific application. This broadly applicable magnetoid model system offers a simple yet powerful tool to enable locally guided mechanobiological studies in organoid model systems in development and disease.

## Materials and Methods

### Human PSC lines

The human PSC lines used in this study are 1) NCRM-1 (RRID:CVCL_1E71) hPSC line from NIH Center for Regenerative Medicine (CRM), Bethesda, USA. 2) ZO1 hPSC line (Mono-allelic mEGFP-Tagged TJP1 WTC, Coriell institute for Medical Research), 3) hPSC F-actin reporter line provided by Catherine Verfaillie, Stem Cell institute Leuven.

### Human PSC culture

Matrigel-coated 6 well plates were used to maintain hPSCs cultured to 60-70% confluence. Human PSC colonies were passaged using a treatment of Dispase (Sigma) for 4 m at 37°C followed by PBS washes. 1 mL of Essential 8 (E8) - Flex Medium Kit (ThermoFisher Scientific) supplemented with 1% Penicillin Streptomycin (GIBCO) and Y-27632 Rock inhibitor (ROCKi) (Hellobio) at 10 μM was used to passage colonies after gentle scraping and agitation with a pipette to break down the colonies. Colonies were passaged 1:6 and incubated in 2 mL of E8-Flex medium supplemented with ROCKi at 10 μM for 24 h. The medium was replaced by 4 mL of fresh E8-Flex and the colonies were incubated for an additional 48 h when they reached 60-70% confluence.

### Magnetic nanoparticle (MNP) preparation and magnetic cluster (MagC) labeling with red fluorescent particles

100 mg of Fe_3_O_4_ magnetic nanoparticles (MNPs) (Sigma-Aldrich, 20-40 nm) were first weighed under sterile conditions and added to 1 mL of E8-Flex media, in a 1.5 mL Eppendorf tube, resulting in a stock MagC mixture with a concentration of 100 mg/mL. The solution was pipetted thoroughly and subsequently sonicated for 5 m to break apart any aggregation. To label the MagCs, 20 μL (2 mg of MagCs) were taken from the stock mixture, immediately following sonication, and added to 2 mL of E8-Flex medium supplemented with 10 μM of ROCKi. The mixture was then sonicated for 3 m to break up aggregation. Next red fluorescent beads (FluoSpheres Carboxylate-Modified Microspheres, 200 nm, Invitrogen) were diluted 1:10 in E8-Flex supplemented with 10 μM of ROCKi and sonicated for 5 m to break up any aggregation. Subsequently, 5 μL of the diluted fluorescent particles were added to the 2 mL MagC-medium mixture to obtain a final composition of 0.025% v/v (1:4000 dilution of the stock solution). The mixture was immediately sonicated for 3 m to increase MNP-fluorescent particle interaction resulting in efficient labeling of the MagCs. This created a 2X magnetic cluster (MagC) solution.

### Magnetization of human induced pluripotent stem cells

Human hPSCs, at a confluence of 60–70%, were washed with PBS three times, followed by the application of 1 mL of TrypLE Express (GIBCO) at 37°C for 3 m for dissociation. Once dissociated into single cells, 9 mL of DMEM/F12 medium supplemented with 20% FBS (GIBCO) was added for neutralization and TrypLE Express washing. The cells were then centrifuged at 300 RCF for 3 m After discarding the supernatant, a second wash was applied by adding 10 mL of DMEM/F12 medium containing 20% FBS followed by centrifugation at 300 RCF for 3 m. The supernatant was discarded and the cells were resuspended in 1 mL E8-Flex medium supplemented with 10 μM of ROCKi. A cell count was performed and the volume of medium was adjusted to obtain cell density of 500,000 cells/mL. A volume of 1 mL was added to a Matrigel-coated well of a 6-well-plate. Immediately after, 1 mL of the 2X MagC solution was added and gently mixed with the cells using a 1,000 μL pipette. The same protocol was followed for the control cells, except the final 1 mL medium addition contained no MagCs. The cells were incubated for 24 h (37°C, 5% CO_2_) to obtain mhPSCs and control hPSCs.

### Magnetoid protocol

A 24-well AggreWell™400 plate (Stemcell Technologies) was first prepared following the manufacturer’s anti-adherence treatment recommendations. Magnetized and control cells were washed, dissociated, and centrifuged following the above protocol. After a cell count, enough volume of dissociated cells was added to 2.5 mL of E8-Flex medium supplemented with 10 μM of ROCKi to obtain a density of 89,100 cells/mL. A similar procedure was followed for the mhPSCs, however, obtaining 1 mL of E8-Flex medium containing 10 μM of ROCKi and a cell density of 900 cells/mL.

Next, the 24-well AggreWell™400 plate was prepared for cell aggregation. First, 1 mL of mhPSCs was added to a well and aggregated immediately at 300 RCF for 3 m. Subsequently, in the same well, 1 mL hPSCs was added dropwise to the liquid-air interface using a 200 μL pipette. In the control well, 1.01 mL of hPSCs was added directly and topped with 0.99 mL of E8-Flex supplemented with 10 μM of ROCKi. The cells were immediately aggregated at 300 RCF for 3 m.

Once aggregated, another standard 24-well-plate was used to house a magnet (Supermagnete, N45, 15 × 15 × 15 mm) with the magnetization axis pointing vertically. The magnet was placed inside one of the wells and covered by the well-plate cover. The well was chosen such that it matched the position the the well containing the mhPSCs in the 24-well AggreWell™400 plate. Next, starting from a vertical position of more than 20 cm directly above the 24-well-plate with the magnet, the AggreWell™ was lowered until it was resting securely above the well-plate containing the magnet. The assembly was then moved to the incubator and the cells were incubated (37°C, 5% CO_2_) for 24 h.

After the incubation period, the assembly was gently moved out of the incubator and the AggreWell™ was lifted in a vertical manner until it was approximately 20 cm away from the well-plate containing the magnet. This ensured the magnetic aggregates were not magnetically disturbed. 1 mL of media was gently taken out. The magnetic and nonmagnetic aggregates were then separately suspended in the remaining 1 mL of medium using a 1 mL pipette (a wide tip bore should be used to avoid damaging the aggregates). The aggregates were then moved to separate 1.5 mL Eppendorf tubes and centrifuged at 150 RCF for 3 m. The supernatant was removed and 12 μL of differentiation medium was added to each aggregate condition.

A 100 μL 2 kPa Polyethylene glycol (PEG) hydrogel was prepared as previously described^14^ for each condition. The 10 μL cell suspension^14^ was replaced by a 10 μL aggregate suspension. 10 μL of aggregate-containing hydrogels were placed in wells of a 96-well-plate. For the magnetically actuated magnetoid condition, the border wells were chosen. For the control and magnetoids without magnetic actuation, a different well-plate was chosen. The well-plates were continuously rotated to prevent organoid aggregation and settling until PEG gelation was visually confirmed^14^. An additional 20 m was required to ensure complete gelation at room temperature conditions. Subsequently 200 μL of differentiation medium (see below) was added to each well. A rectangular magnet (Supermagnete, N45, 30 × 30 × 15 mm) was placed next to the well-plate containing the magnetoid for actuation. The magnet was placed on a 1.5 mm plastic spacer so that it was not directly touching the incubator shelf. The well-plate containing the control and non-actuated magnetoids was placed at least 20 cm away from the magnet to reduce magnetic disturbance.

### Human neural tube differentiation media

Organoids were treated immediately after PEG embedding for 3 days with neural differentiation medium comprised of a 1:1 mixture of neurobasal medium (GIBCO) and DMEM/F12 (GIBCO), 1% N2 (GIBCO), 2% B-27 (GIBCO), 1 mM sodium pyruvate MEM (GIBCO), 1 mM glutamax (GIBCO), 1 mM non-essential amino acids (GIBCO) and 2% Penicillin Streptomycin (GIBCO) and supplemented with 10 μM of ROCKi. To induce floor plate identity, organoids were treated from days 3 - 5 with ROCKi-free neural differentiation medium supplemented with retinoic acid (RA) (Stemcell Technologies) at 0.25 μM and smoothened agonist (SAG) (Stemcell Technologies) at 1 μM. The organoids were subsequently treated with ROCKi-free neural differentiation medium until end point day 11 and the medium was fully refreshed every 2 days.

### Human paraxial mesoderm differentiation media

Organoids were treated immediately after PEG embedding for 3 days with paraxial mesoderm differentiation medium comprised of DMEM/F12 (GIBCO), 1% N2 (GIBCO), 2% B-27 (GIBCO), 1 mM sodium pyruvate MEM (GIBCO), 1 mM glutamax (GIBCO), 1 mM non-essential amino acids (GIBCO) and 2% Penicillin Streptomycin (GIBCO) and supplemented with 10 μM of ROCKi, 10 μM of CHIR99021 (Tocris) and 500 nM of LDN193189 (Stemgent). Next the organoids were treated from days 3 - 6 with ROCKi-free mesoderm differentiation medium supplemented with 10 μM of CHIR99021, 500 nM of LDN193189, and 20 ng/mL of fibroblast growth factor 2 (FGF2) (Peprotech). Finally the organoids were treated from days 6 - 8 with ROCKi-free mesoderm differentiation medium supplemented with 500 nM LDN193189, 20 ng/mL FGF2, 2 ng/mL insulin-like growth factor 1 (IGF1) (Peprotech), and 10 ng/mL hepatocyte growth factor (HGF) (Peprotech).

### Magnetic nanoparticle toxicity study

Fluorescently labeled MagCs were prepared following the MagC labeling protocol to obtain 2X concentration solutions of 20, 200, 2,000, and 10,000 µg/mL. The respective fluorescent particle concentration for each condition was adjusted to maintain their ratio to MNPs. Single hPSCs were plated in wells of a 6-well-plate at a density of 100,000 cells/mL in a volume of 1 mL of E8-Flex supplemented with 10 μM ROCKi. Next 1 mL of the 2X MagC solutions was added to the respective wells and mixed gently with a 1,000 μL pipette. The cells were incubated for 24 h, at which point the medium was replaced with ROCKi-free E8-Flex medium and incubated for an additional 48 h. Media change was done carefully to not disturb the MagCs in the wells. Cells were then washed, dissociated, centrifuged and resuspended in 1 mL of E8-Flex medium supplement with 10 μM of ROCKi. A cell count was performed and the total amount of cells per condition was evaluated and normalized to that of the control condition. These values were reported to compare the toxicity of MagCs on hPCS. In separate wells with intact colonies, Calcien AM (Thermofisher) was used to visualize live cells in control and magnetized conditions. The cells were then fixed in paraformaldehyde (PFA 4%, Sigma-Aldrich) for 2 h and gently washed with PBS. The cells were then treated with a permeabilization and blocking solution comprised of 0.3% Triton X (PanREAC AppliChem) and 0.5% BSA (Sigma–Aldrich) in PBS for 30 m. Hoechst was then used to visualize DNA and applied at a concentration of 1:2000 in permeabilization and blocking solution for 2 h at 4°C followed by 3 PBS washes. Representative images were then obtained using an inverted microscope (Zeiss Axio Observer Z1; Carl Zeiss MicroImaging) equipped with a Colibri LED light source and a 10X air objective.

### Magnetic cluster localization

ZO1 hPSCs were magnetized using a 1 mg/mL MagC solution following the magnetization protocol. The cells were then imaged using confocal microscopy (Leica SP8 DIVE, Leica Microsystems). 3-D reconstructed images were generated from image stacks to demonstrate the localization of MagCs and ZO1 (apical).

### Magnetic cluster size analysis

MagCs were prepared and labeled as described in the above protocol to yield a 2 mg/mL 2X-2 mL solution. A volume of 5 μL was taken and added to 1 mL of DMEM medium and sonicated for 3 m. A 100 μL droplet of the mixture was taken and added to a glass slide. The droplet was imaged using an inverted microscope (Zeiss Axio Observer Z1; Carl Zeiss MicroImaging) equipped with a Colibri LED light source and a x40 air objective. MagCs sizes were assessed using the particle analysis function in ImageJ.

### Magnetization efficiency

Magnetized hPSCs were prepared with final MagC concentrations of 10, 100 and 1,000 μm/mL following the magnetization protocol. After 24 h incubation, the cells were washed, dissociated, centrifuged and resuspended in 1 mL of E8-Flex medium supplement with 10 μM ROCKi. A 100 μL droplet of the mixture was taken and added to a glass slide. The glass slide was placed under an inverted microscope (EVOS, Invitrogen) and visualized using a x4 objective. An image was acquired showing the cell positions as they settled towards the glass slide. A magnet (Supermagnete, N45, 15 × 15 × 15 mm) was placed at a distance not greater than 1 mm from the droplet edge. After 2 m, a second image was taken of the same droplet and position. Using ImageJ, a visual comparison determined the percentage of cells that underwent magnetophoresis. Two other conditions were also assessed, 1) control unmagnetized hPSCs, and 2) hPSCs that had been instantly mixed with a final MagC concentration of 1,000 μg/mL, following magnetization protocol.

### Magnetic simulations (FEMM) and force estimation

The program Finite Element Method Magnetics (FEMM, version 4.2) was used to conduct two-dimensional axisymmetric magnetics simulations to compute the magnetic field produced by the various magnets used in the study. First the profile area of the used magnet was constructed and assigned a NdFeB N45 material property from the built-in FEMM material library. The well-plate, the PEG hydrogel and the organoid tissue were all considered permeable to the magnetic field and thus ignored in the simulation. All unfilled spaces around the magnet was assigned the material properties of air from the built-in FEMM material library. Standard meshing was chosen prior to running the simulation.

To evaluate the magnetic force produced by MagCs in magnetoids plated in a hydrogel in a 96-well-plate, a straight line was taken from the corner of the magnet and extended outwards into the air environment. The line length spanned the entire length of a well-plate and the magnetic field values, **B**, were extracted in Tesla. Next, the gradient of the squared field, ∂|**B**|^2^/∂y, was calculated along the length of the line section. The magnetic force density, 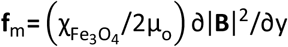, was then calculated where the magnetic volume susceptibility of Fe_3_O_4_ was assumed for MNPs as 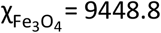 and the permeability of free space μ_o_ =4π×10^−7^. Each well directly facing the magnet (e.g. six adjacent wells that form a column on a 96-well-plate) was divided into six horizontal tranches with equal thickness. Each tranche was assigned one **f**_m_ value using the simulation data and based on the distance separating the centroid of the tranche to the magnet surface. For any magnetoid in a tranche, the magnetic force 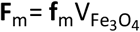, where 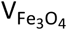 was the volume of the MagC. In this way, the force generated by any MagC in any magnetoid located at any distance away from the magnet surface could be estimated. The MagC volume was approximated as a cylinder of volume 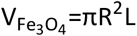, where R is half the length of the short axis of the MagC projected area, and L is the length of long axis of the MagC projected area as measured on ImageJ.

### Magnetic force and single cell fate correlation

The magnetic force was measured as detailed above. Next the distance that separates the centroid of FOXA2+ and PAX6+ nuclei and that of the MagC in the same organoid was measured. Both the distance and force were then reported in a correlation plot (**Supplementary Fig. 4**). Only distances toward the biased zones for each fate (FOXA2 – towards the magnet, PAX6 – away from the magnet) was reported.

### Magnetic force analysis on single cells

Magnetized hPSCs were prepared with a final MagC concentration of 1,000 μm/mL following the magnetization protocol. After 24 h incubation, the cells were washed, dissociated, counted, centrifuged, and resuspended in E8-Flex medium supplement with 10 μM ROCKi with a volume to obtain ∼10-100 cell/μL. This low concentration reduced cell-cell flow perturbation. A glass slide was prepared where a magnet (Supermagnete, N45, 3 × 3 × 3 mm) was glued in place with the magnetization axis aligned with the slide’s long axis. The glass slide was then mounted onto the stage of an inverted microscope (Zeiss Axio Observer Z1; Carl Zeiss MicroImaging, x5 objective). Next, 100 µL of the cell suspension was placed onto a glass slide. As the cells underwent magnetophoresis, they began to move through the medium toward the magnet surface and a video (5 frames per second) using the brightfield channel was taken. Using ImageJ, the distance (measured between forward cell edges in consecutive frames) traveled for each cell between frames was measured and divided by 0.2 s to obtain a velocity **u** (∼100 µm/s). For a cell radius R of ∼10 µm, this velocity resulted in a Reynold’s number ≪ 1, suggesting Stoke’s flow conditions where **F**_D_ = **F**_M_ where **F**_D_ is the drag force and **F**_M_ is the magnetic force acting on the MagCs. For each cell, **F**_D_=6πμR**u**, where µ = 8.9 · 10-4 Pa⋅s is the dynamic viscosity of water, R is the cell radius assuming a spherical geometry, **u** is the calculated cell velocity between frames. The magnetic field produced by the 3 × 3 × 3 mm magnet was then simulated and a force density plot was generated. Knowing **F**_M_ and using the force density plot we could estimate the volume of MagCs attached the cells ∼7.3 ×10^3^ µm^3^. When comparing this value to those measured for MagCs in magnetpoid, we saw a difference, suggesting that in addition to the assumed ∼1 mhPSCs that is integrated per magnetoid for the 1% mhPSC condition, other unbound MagCs can also find their way into the cellular construct during the aggregation step. It is therefore more accurate, for the purpose of evaluating forces in magnetoids, to estimate the MagC volume in magnetoids directly from images and not simply assume a volume based on the single cell **F**_M_ measurements. These single cell magnetophresis experiments infered proper attachment of the MagCs to the cells.

### Short and long term magnetoid actuation

A short term magnetic actuation was conducted on magnetoids embedded in 2 kPa PEG hydrogels, prepared as discussed above and plated in a 96 well plate. Day 11 hNTmOs were observed using an inverted microscope (EVOS, Invitrogen). A magnet (Supermagnete, N45, 15 × 15 × 15 mm) was placed at a ∼45° angle (directly next to the adjacent well) with the magnetization axis perpendicular to the plate edge. This setup still allowed MagCs to reorient toward the magnet. An image was then taken of the magnetoid every 5 s for 30 s. Using ImageJ, the displacement of the tissue around the MagC as it reorients towards the magnet was traced and measured by choosing a visible and trackable tissue feature on the brightfield channel. We measured the displacements caused by whole magnetoid rotation and subtracted these from the tissue displacements. The field of view was divided into an array of tissue sections and the displacements, d_tissue_, in each tissue section were normalized to the displacement of the part of the MagC, d_M_, in that row to provide a dimensionless displacement value d*. The distance, x, from the centroid of each tissue section to the edge of the MagC in the same row was normalized to the displacement of the part of the MagC in that row to provide a dimensionless coordinate value c*. A d*-c* correlation and heatmap was then reported.

### Cytoskeleton rearrangement study

A similar setup as the one discussed above was used. However, here the magnet was placed for longer durations of 5, 30, 60, and 120 m under incubation conditions of previously magnetically actuated day 11 hNTmOs. At time t_0_ without the magnet placement, the reference angle of the MagC was imaged. At every subsequent time point, the magnet was taken away and the new MagC angle was imaged. The recoil angle, α, was calculated by subtracting the reference angle from the new angle and was then reported.

To observe cytoskeletal rearrangement, the F-actin hPSC reporter line was used along with the above setup, however, using a confocal microscope (Leica SP8 DIVE, Leica Microsystems) with incubation capability (37°C, 5% CO_2_). Here, the magnet remained in place and an image was taken to report the cytoskeleton architecture of the magnetoids at every time point of 5, 30, 60 and 120 m. Next, using ImageJ, intensities of F-actin were observed on the tension side of the MagC as well as on the directly opposite pole of the magnetoid. The intensities were then normalized to intensities at time t_5 m_. Due to the setup, a time t_0 m_ image was not possible without disrupting the configuration, and thus was avoided.

### Growth and proliferation studies

To determine the size of hNTmOs, day 11 magnetoids were imaged using an inverted microscope (EVOS, Invitrogen). Using ImageJ, the projected area of organoids was traced and converted to an equivalent diameter assuming a circular area.

To determine the biased growth of magnetoids when compared to control or unactuated magnetoids, magnetoids were imaged at days 1 and 11. A contour line was traced at days 1 and 11 and the images were superimposed. Next, a line section, P_m_, was traced connecting the closest point to the magnet surface of the contours at days 1 and 11. A similar line section, P_a_, was traced to connect the furthest point to the magnet at both days. The ratio of the growth towards the magnet, P_m_, and growth away from the magnet, P_a_, was calculated where a value <1 and >1 indicated biased growth away and toward the magnet respectively while a value ∼1 represented an unbiased growth direction.

To determine the proliferation bias, day 11 hNTmOs were fixed and stained with EdU and images were taken using an inverted microscope (Zeiss Axio Observer Z1; Carl Zeiss MicroImaging, x10 objective). In the case of actuated magnetoids, a line segment dividing the magnetoids into two halves and passing through the MagC centroid was traced. For the control organoids and unactuated magnetoids, the line segment passed through the organoid centroid. The average EdU intensity in the halves facing the magnet, EdU_m_, was normalized to that of the respective halves facing away from the magnet, EdU_a_, where a value <1 and >1 indicated biased proliferation away and toward the magnet respectively while a value ∼1 represented an unbiased proliferation direction.

### Fate specification and patterning studies

The induction efficiencies of FOXA2 and PAX6 were visually inspected across the various conditions using and inverted microscope (Zeiss Axio Observer Z1; Carl Zeiss MicroImaging, 10X objective). Pattering efficiencies of FOXA2 and PAX6 in organoids were assessed as previously reported^14^. Briefly, fluorescent images were assessed using ImageJ to obtain the area of the expression region of FOXA2 or PAX6 as well as the average intensity of the expression in that area. Next, the projected area of the entire organoid or magnetoid was evaluated and an average intensity of the fate expression was obtained. We next evaluated the area ratio (AR) where the fate expression area was divided by the organoid area. We also evaluate the intensity ratio (IR) where the average fate intensity of the expression region was divided by that of the organoid or magnetoid area. The fate was considered as scattered for (1) AR > 0.5, and (2) AR < 0.5 but IR < 20%. By contrast, the fate expression was found to be pattered for AR < 0.5 and IR > 20%.

To evaluate fate directional bias upon internal force generation, inverted microscopy images were assessed in a similar manner as the EdU stains, however only for actuated magnetoids and for FOXA2 and PAX6 separately.

For the magnetic paraxial mesoderm organoids, the hPSC-PAX3 reporter line was used and the induction efficiency was reported along the 8 day experimental timeline using an inverted microscope (EVOS, Invitrogen). The PAX3 directional bias upon internal force generation was then evaluated in similar manner as the EdU, FOXA2, and PAX6 expressions.

### Quantification and statistical analysis

Two-way ANOVA statistical tests and unpaired two-tailed t-test with corrections were used where appropriate with a 95% confidence interval (GraphPad Prism 6, Version 6.01, GraphPad Software, Inc.). Statistical significance was considered for all comparisons with p < 0.05.

## Acknowledgement

This work was supported by the FWO grant G087018N and FWO postdoctoral fellowship 1217220N, Interreg Biomat-on-Chip grant and Vlaams-Brabant and Flemish Government co-financing, KU Leuven grants C14/17/111andC32/17/027and King Baudouin Foundation grant J1810950-207421.

## Supplementary Figures

**Supplementary Fig.1:**
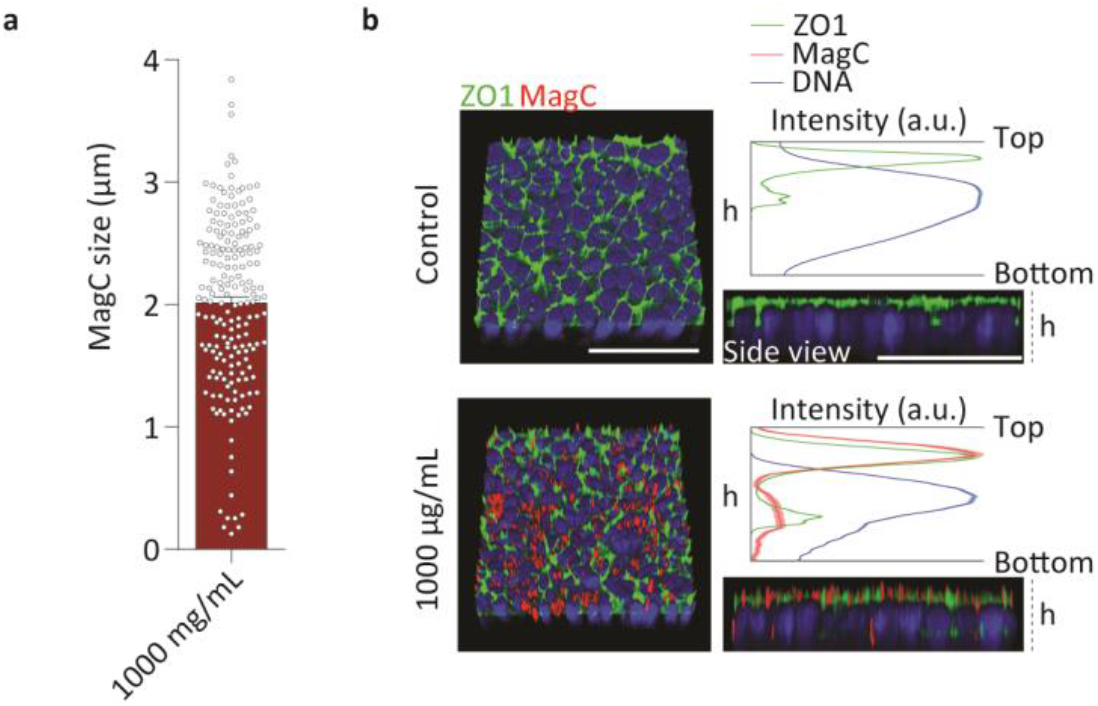
**a** MagC sizes after fluorescent tagging (n = 3 for >200 MagCs). **b** 3-D reconstructed representative images showing ZO1 and MagC localization in control and MagC (5,000 μg/mL) conditions (n = 3). Profiles showing ZO1, MagC and nuclei vertical position (n = 3). Scalebar 50 μm. Error bar SEM.

**Supplementary Fig.2:**
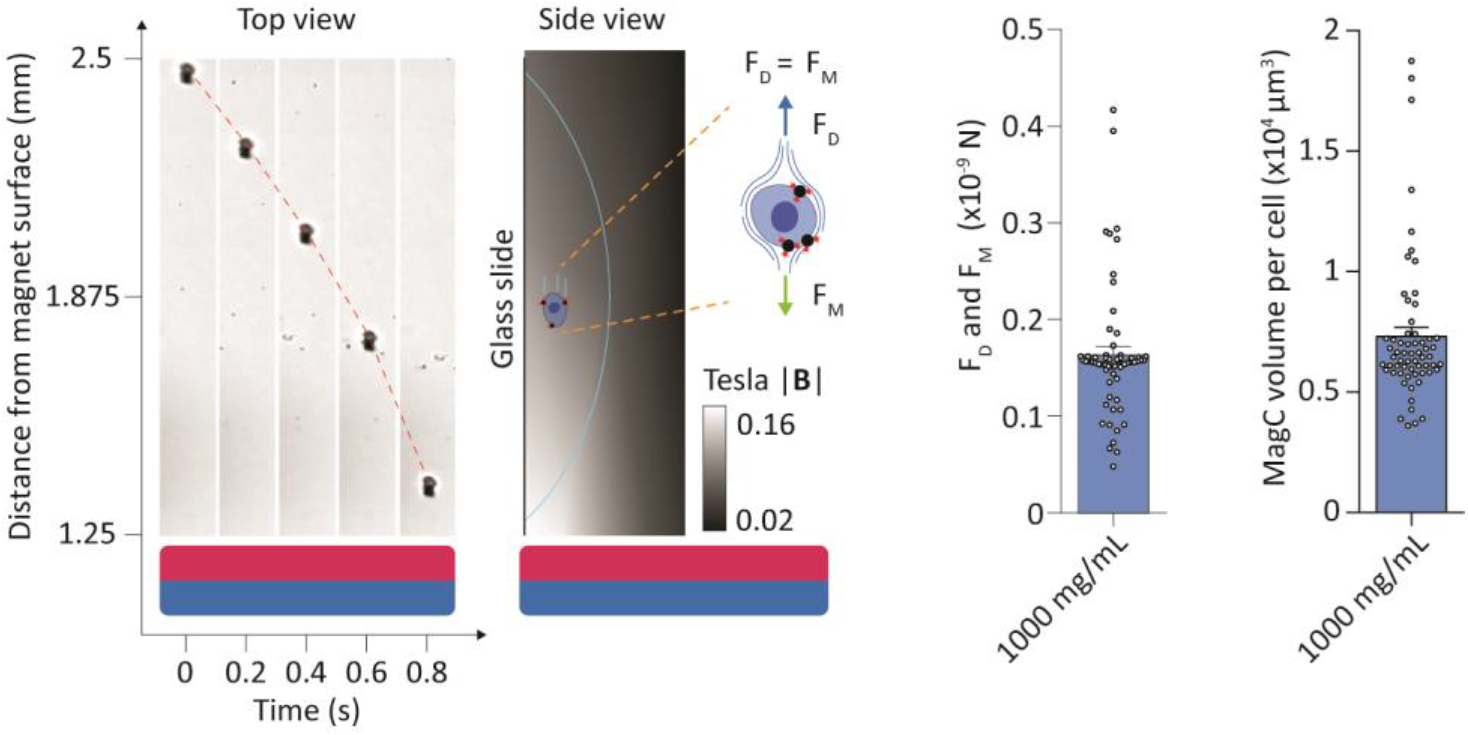
Representative images of a mhPSC undergoing magnetophoresis. Corresponding magnetic field as simulated by a 3 × 3 × 3 mm N45 magnet (n = 3). Schematic representation of Stoke’s drag and magnetic forces. Stoke’s drag and magnetic forces on mhPSCs undergoing magnetophoresis and corresponding MagC volumes (n = 3 for >50 cells). Error bar SEM.

**Supplementary Fig.3:**
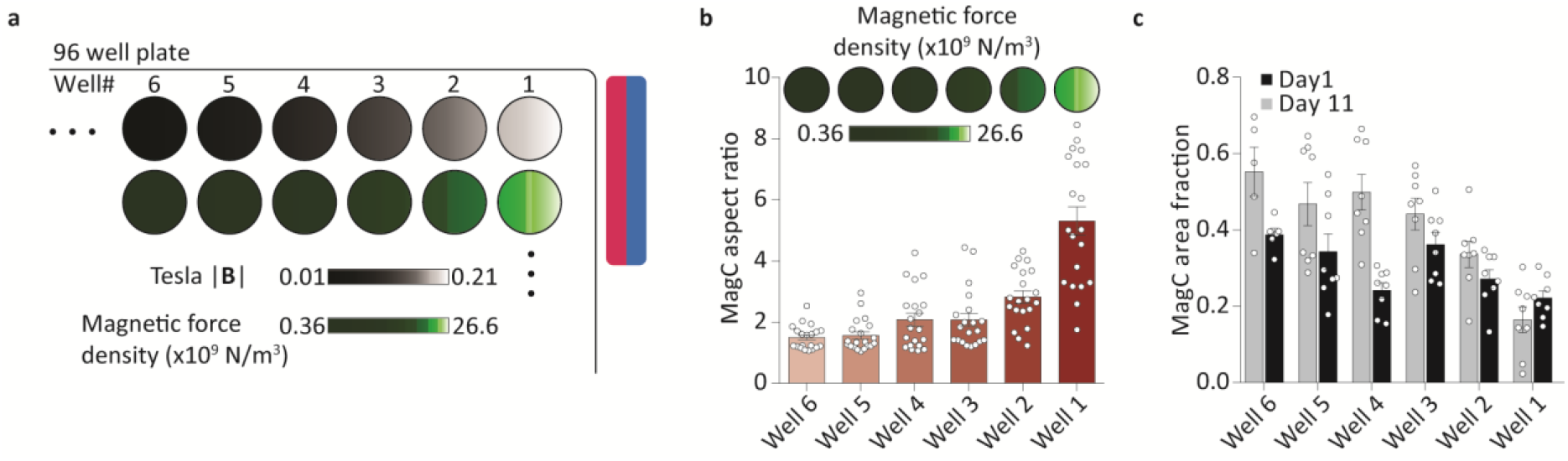
**a** Magnetic field and force densities in wells of a 96 well-plate at various distances from a magnet surface as simulated using a 30 × 30 × 15 mm N45 permanent magnet. **b** MagC aspect ratio of day 11 hNTmOs in wells of a 96 well-plate at various distances from the magnet surface (n = 2 for a total of 20 hNTmOs per well) **c** MagC area ratio of day 1 and 11 hNTmOs in wells of a 96 well-plate at various distances from the magnet surface (n = 1 for a total of 8 hNTmOs per well except for well 6 (day 1 5 hNTmOs and day 11 4 hNTmOs)). Error bar SEM.

**Supplementary Fig.3:**
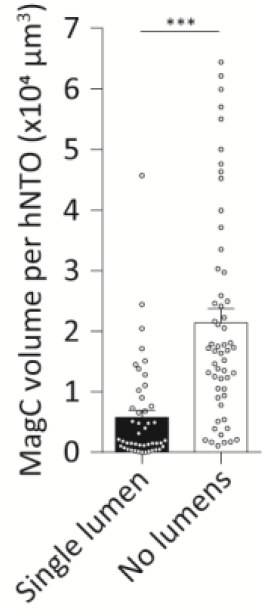
Estimation of MagC volumes in day 11 hNTmOs with single or no lumen formation (n = 3 for >50 magnetoid per condition). Error bar SEM.

**Supplementary Fig.4:**
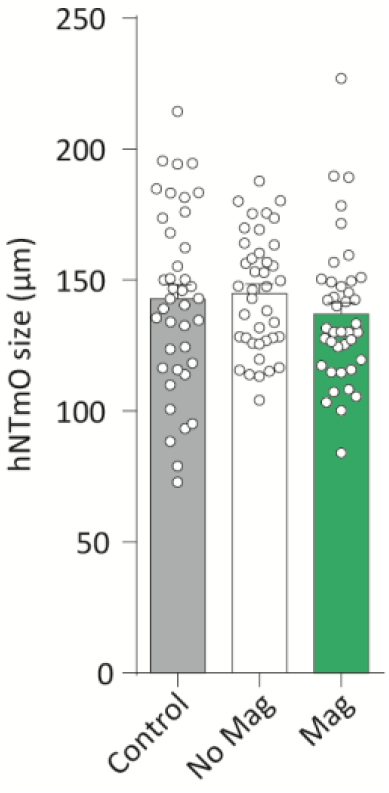
Estimation of MagC volumes in day 11 hNTmOs with single or no lumen formation (n = 3 for a total of 40 magnetoids per condition). Error bar SEM.

**Supplementary Fig.5:**
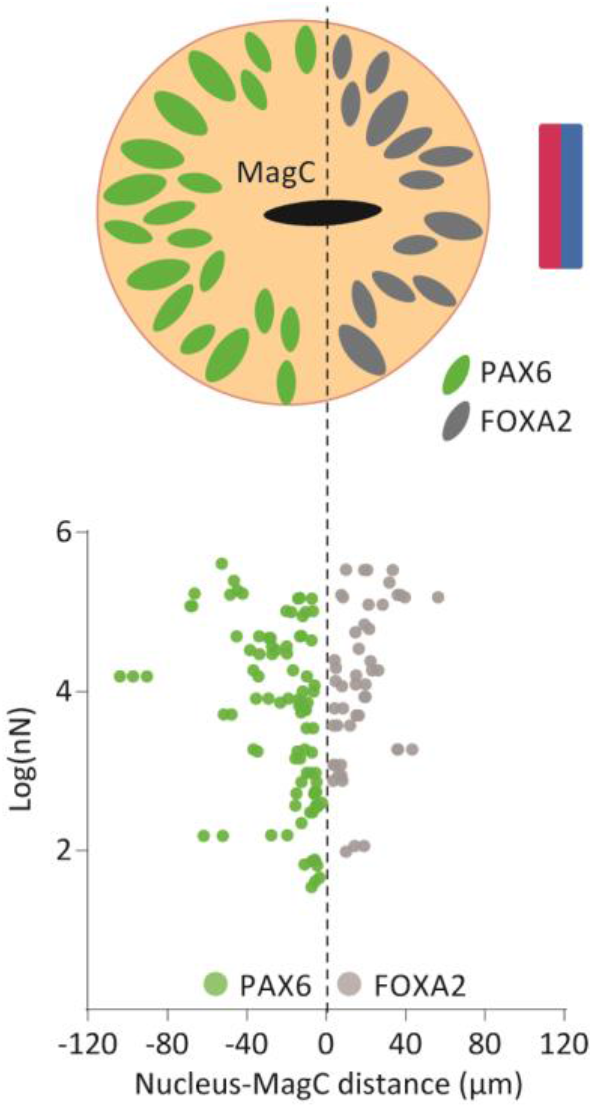
Schematic representation of biased FOXA2 and PAX6 expressions under magnetic actuation condition. Correlation of the magnetic log(nN) force generated by a MagC and the distance separating the MagC and the nucleus expressing FOXA2 or PAX6 fates (n = 4, data from **Fig.4c**).

**Supplementary Fig.7:**
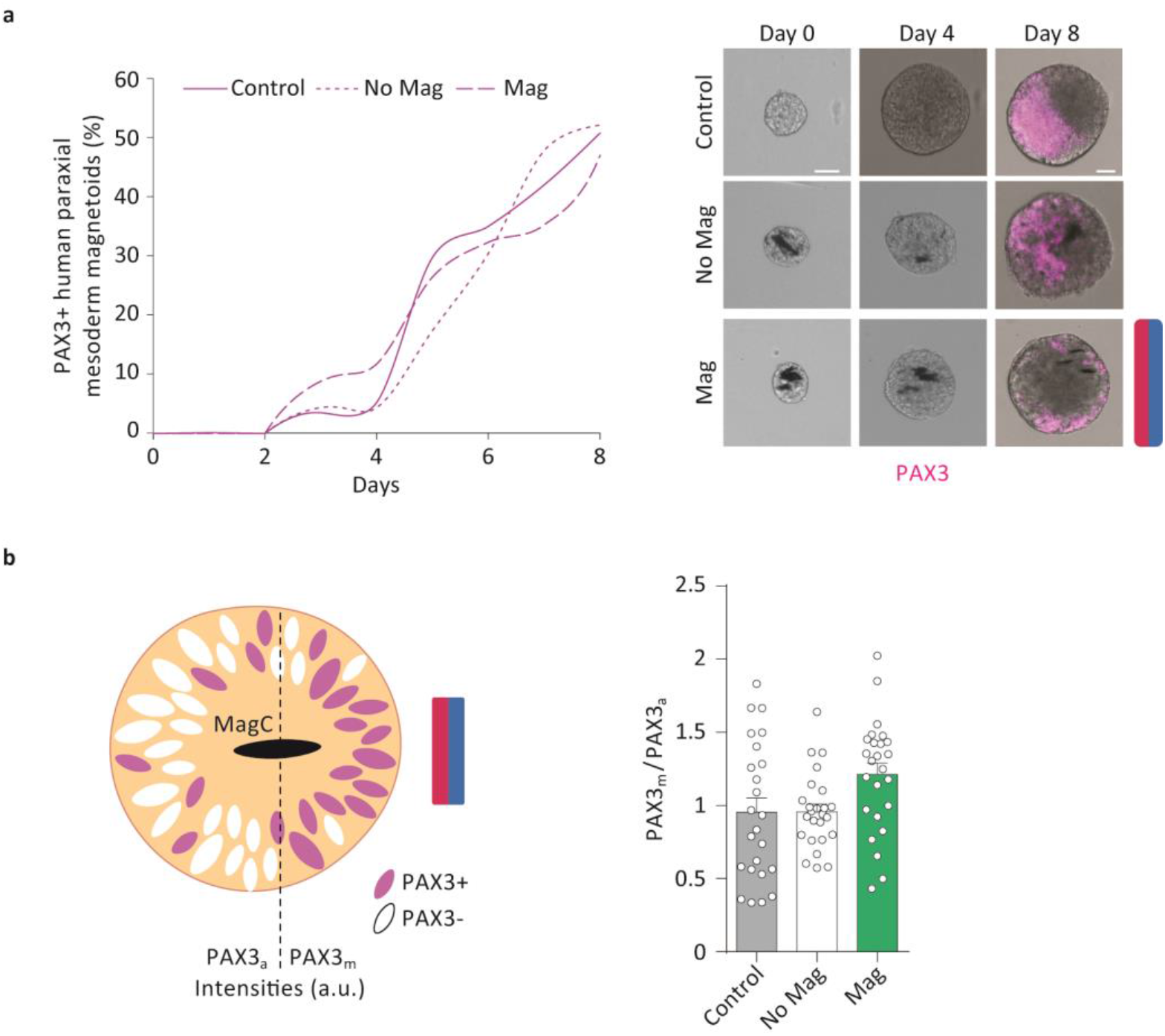
**a** PAX3 expression in human paraxial mesoderm magnetoids under various conditions (n = 1 for a total of 25 magnetoids for all data points per condition). Representative images of human paraxial mesoderm magnetoids under various conditions (n = 1 for a total of 25 magnetoids). **b** Schematic representation of biased PAX3 expression under magnetic actuation condition. Quantification of PAX3 patterning in hNTmOs under various actuation condition (n = 1 for a total of 25 magnetoids per condition). Scalebar 50 μm. Error bar SEM.

## References

1. Heisenberg, C. P. & Bellaïche, Y. XForces in tissue morphogenesis and patterning. Cell 153, 948 (2013).

2. LeGoff, L. & Lecuit, T. Mechanical forces and growth in animal tissues. Cold Spring Harb. Perspect. Biol. 8, 1–17 (2016).

3. Davidson, L. A. Mechanical design in embryos: Mechanical signalling, robustness and developmental defects. Philos. Trans. R. Soc. B Biol. Sci. 372, (2017).

4. Dupont, S. et al. Role of YAP/TAZ in mechanotransduction. Nature 474, 179–184 (2011).

5. Chan, C. J., Heisenberg, C. P. & Hiiragi, T. Coordination of Morphogenesis and Cell-Fate Specification in Development. Curr. Biol. 27, R1024–R1035 (2017).

6. Glaser, O. C. On the mechanism of morphological differentiation in the nervous system. I. The transformation of a neural plate into a neural tube. Anat. Rec. 8, 525–551 (1914).

7. Brown, M. G., Hamburger, V. & Schmitt, F. O. Density studies on amphibian embryos with special reference to the mechanism of organizer action. J. Exp. Zool. 88, 353–372 (1941).

8. Nishimura, T., Honda, H. & Takeichi, M. Planar cell polarity links axes of spatial dynamics in neural-tube closure. Cell 149, 1084–1097 (2012).

9. Schoenwolf, G. C. & Franks, M. V. Quantitative analyses of changes in cell shapes during bending of the avian neural plate. Dev. Biol. 105, 257–272 (1984).

10. Smith, J. L. & Schoenwolf, G. C. Notochordal induction of cell wedging in the chick neural plate and its role in neural tube formation. J. Exp. Zool. 250, 49–62 (1989).

11. Gjorevski, N. et al. Designer matrices for intestinal stem cell and organoid culture. Nature 539, 560–564 (2016).

12. Cruz-Acuña, R. et al. Synthetic hydrogels for human intestinal organoid generation and colonic wound repair. Nat. Cell Biol. 19, 1326–1335 (2017).

13. Ranga, A. et al. Correction for Ranga et al., Neural tube morphogenesis in synthetic 3D microenvironments. Proc. Natl. Acad. Sci. 114, E3163–E3163 (2017).

14. Abdel Fattah, A. R. et al. Actuation enhances patterning in human neural tube organoids. Nat. Commun. 12, 1–13 (2021).

15. Falleroni, F., Torre, V. & Cojoc, D. Cell mechanotransduction with piconewton forces applied by optical tweezers. Front. Cell. Neurosci. 12, 1–11 (2018).

16. Zamir, E. A. & Taber, L. A. Material Properties and Residual Stress in the Stage 12 Chick Heart During Cardiac Looping. J. Biomech. Eng. 126, 823–830 (2005).

17. Marrese, M. et al. Investigating the Effects of Mechanical Stimulation on Retinal Ganglion Cell Spontaneous Spiking Activity. Frontiers in Neuroscience 13, (2019).

18. Kim, H. Y., Jackson, T. R., Stuckenholz, C. & Davidson, L. A. Tissue mechanics drives regeneration of a mucociliated epidermis on the surface of Xenopus embryonic aggregates. Nat. Commun. 11, 1–10 (2020).

19. Serwane, F. et al. In vivo quantification of spatially varying mechanical properties in developing tissues. Nat. Methods 14, 181–186 (2017).

20. Tseng, P., Judy, J. W. & Di Carlo, D. Magnetic nanoparticle-mediated massively parallel mechanical modulation of single-cell behavior. Nat. Methods 9, 1113–1119 (2012).

21. Du, V. et al. A 3D magnetic tissue stretcher for remote mechanical control of embryonic stem cell differentiation. Nat. Commun. 8, (2017).

22. Panariti, A., Miserocchi, G. & Rivolta, I. The effect of nanoparticle uptake on cellular behavior: Disrupting or enabling functions? Nanotechnol. Sci. Appl. 5, 87–100 (2012).

23. Schweiger, C. et al. Quantification of the internalization patterns of superparamagnetic iron oxide nanoparticles with opposite charge. J. Nanobiotechnology 10, 1–11 (2012).

24. Hoskins, C., Cuschieri, A. & Wang, L. The cytotoxicity of polycationic iron oxide nanoparticles: Common endpoint assays and alternative approaches for improved understanding of cellular response mechanism. J. Nanobiotechnology 10, (2012).

25. Liu, X. et al. A brief review of cytotoxicity of nanoparticles on mesenchymal stem cells in regenerative medicine. Int. J. Nanomedicine 14, 3875–3892 (2019).

26. Rahman, A., Fattah, A. & Ranga, A. Nanoparticles as Versatile Tools for Mechanotransduction in Tissues and Organoids. 8, 1–12 (2020).

27. Selman, G. G. The Forces Producing Neural Closure in Amphibia. Development 6, 448–465 (1958).

28. Deng, L., Fairbank, N. J., Fabry, B., Smith, P. G. & Maksym, G. N. Localized mechanical stress induces time-dependent actin cytoskeletal remodeling and stiffening in cultured airway smooth muscle cells. Am. J. Physiol. Cell Physiol. 287, C440–8 (2004).

29. Yamada, T., Placzek, M., Tanaka, H., Dodd, J. & Jessell, T. M. Control of cell pattern in the developing nervous system: Polarizing activity of the floor plate and notochord. Cell 64, 635–647 (1991).

30. Molina, A. & Pituello, F. Playing with the cell cycle to build the spinal cord. Dev. Biol. 432, 14–23 (2017).

31. Bonnet, F. et al. Neurogenic decisions require a cell cycle independent function of the CDC25B phosphatase. Elife 7, e32937 (2018).

32. Meinhardt, A. et al. 3D reconstitution of the patterned neural tube from embryonic stem cells. Stem Cell Reports 3, 987–999 (2014).

33. Zagorski, M. et al. Decoding ofposition in the developing neural tube from antiparallel morphogen gradients. Science (80-.). 356, 1–5 (2017).

34. Chal, J. et al. Recapitulating early development of mouse musculoskeletal precursors of the paraxial mesoderm in vitro. Development 145, dev157339 (2018).

35. Xue, X. et al. Mechanics-guided embryonic patterning of neuroectoderm tissue from human pluripotent stem cells. Nat. Mater. 17, 633–641 (2018).

